# Pbp1 stabilizes and promotes the translation of Puf3-target mRNAs involved in mitochondrial biogenesis

**DOI:** 10.1101/2022.09.14.507933

**Authors:** Floortje van de Poll, Benjamin M. Sutter, Chien-Der Lee, Yu-San Yang, Benjamin P. Tu

## Abstract

Pbp1 (poly(A)-binding protein - binding protein 1) is a cytoplasmic stress granule marker that is capable of forming condensates that function in the negative regulation of TORC1 signaling under respiratory conditions. How mutations in its mammalian ortholog ataxin-2 are linked to neurodegenerative conditions remains unclear. Here, we show that loss of Pbp1 leads to decreases in amounts of mitochondrial proteins whose encoding mRNAs are targets of Puf3, a member of the PUF (Pumilio and FBF) family of RNA-binding proteins. We found that Pbp1 stabilizes and promotes the translation of Puf3-target mRNAs in respiratory conditions, such as those involved in the assembly of cytochrome c oxidase. We further show that Pbp1 and Puf3 interact through their respective low complexity domains, which is required for Puf3-target mRNA stabilization and translation. Our findings reveal a key role for Pbp1-containing assemblies in enabling the translation of mRNAs critical for mitochondrial biogenesis and respiration. They may further begin to explain prior associations of Pbp1/ataxin-2 with RNA, stress granule biology, mitochondrial function, and neuronal health.

## Introduction

Yeast cells are capable of rapidly adapting their metabolism to changes in environmental conditions. When grown in glucose media, yeast cells use glycolysis for energy production and suppress mitochondrial biogenesis. However, in the presence of a non-fermentable carbon source, such as lactate, yeast cells adapt by inducing mitochondrial biogenesis to increase ATP production by oxidative phosphorylation (OXPHOS). The majority of the protein components of the electron transport chain are nuclear-encoded and need to be imported into the mitochondria. This essential process also requires cytosolic protein participants and is tightly regulated according to the cell’s metabolic needs (1).

Puf3 is one such cytosolic protein that plays a key role in the post-transcriptional regulation of mitochondrial biogenesis. Puf3 is a member of the PUF protein family, which are conserved RNA-binding proteins that recognize a consensus motif in the 3’ UTR of their target mRNAs (2). Most of the target mRNAs associated with Puf3 are important for OXPHOS, many of which encode for subunits of mitochondrial ribosomes or respiratory complexes (3–8). Depending on glucose availability, Puf3 can switch the fate of its target mRNAs from decay to translation (9). In the presence of non-fermentable sugars that require respiratory metabolism, Puf3 becomes heavily phosphorylated in its N-terminal low complexity region, leading to the stabilization and translation of its bound transcripts. Furthermore, Puf3 has been found in ribonucleoprotein (RNP) granules (9).

The yeast ortholog of ataxin-2, poly(A) binding protein-binding protein 1 (Pbp1), associates with poly(A)-tails and is also found in RNP granules (10,11). Prior evidence also suggests that Pbp1 is important for the maintenance of mitochondrial function. For instance, Pbp1 overexpression has been shown to rescue various mitochondrial abnormalities including defects in intron splicing, protein import, and mitochondrial damage and death caused by a mutation in the mitochondrial ADP/ATP carrier AAC2 that is equivalent to one in humans with progressive external ophthalmoplegia (12,13). We also demonstrated that Pbp1 negatively regulates TORC1 leading to autophagy and mitophagy in minimal growth conditions that require respiration (14). In addition, loss of Pbp1 can lead to mitochondrial dysfunction, increased sensitivity to oxidative stress, and cell death (14,15). These findings collectively suggest that Pbp1 plays an important role in promoting mitochondrial function.

Here we utilized *S. cerevisiae* to discover a functional relationship between Pbp1, mitochondria, and Puf3. We find that Pbp1 and Puf3 interact through their low-complexity domains and that this interaction facilitates the translation of mRNAs that are targets of Puf3 under respiratory conditions. These findings reveal that Pbp1 supports mitochondrial function via assemblies involved in mRNA translation, which may provide key insights into how ataxin-2 mutations cause neurodegenerative diseases.

## Results

To interrogate a possible role for Pbp1 in mitochondrial function, we assayed mitochondrial protein abundance in response to switching from glycolytic (YPD) to respiratory (YPL) media, conditions that demand mitochondrial biogenesis, in wild type (WT) and *pbp1Δ* cells. We assessed Por1 (mitochondrial porin) and Cox2 (subunit II of cytochrome c oxidase) protein levels using readily available antibodies by immunoblot in YPD and at several time points after switching to YPL (Figure 1A). Por1 protein levels increased over time during growth in respiratory conditions in both WT and *pbp1Δ* cells. Strikingly, Cox2 protein levels were severely decreased in *pbp1Δ* compared to WT cells at all time points. Moreover, *pbp1Δ* cells showed a significantly reduced growth rate in YPL compared to WT cells (Figure 1B), but their growth rate in YPD was normal. These observations are consistent with *pbp1Δ* cells having compromised mitochondrial function due to reduced amounts of proteins required for respiration, such as Cox2.

**Figure 1.**
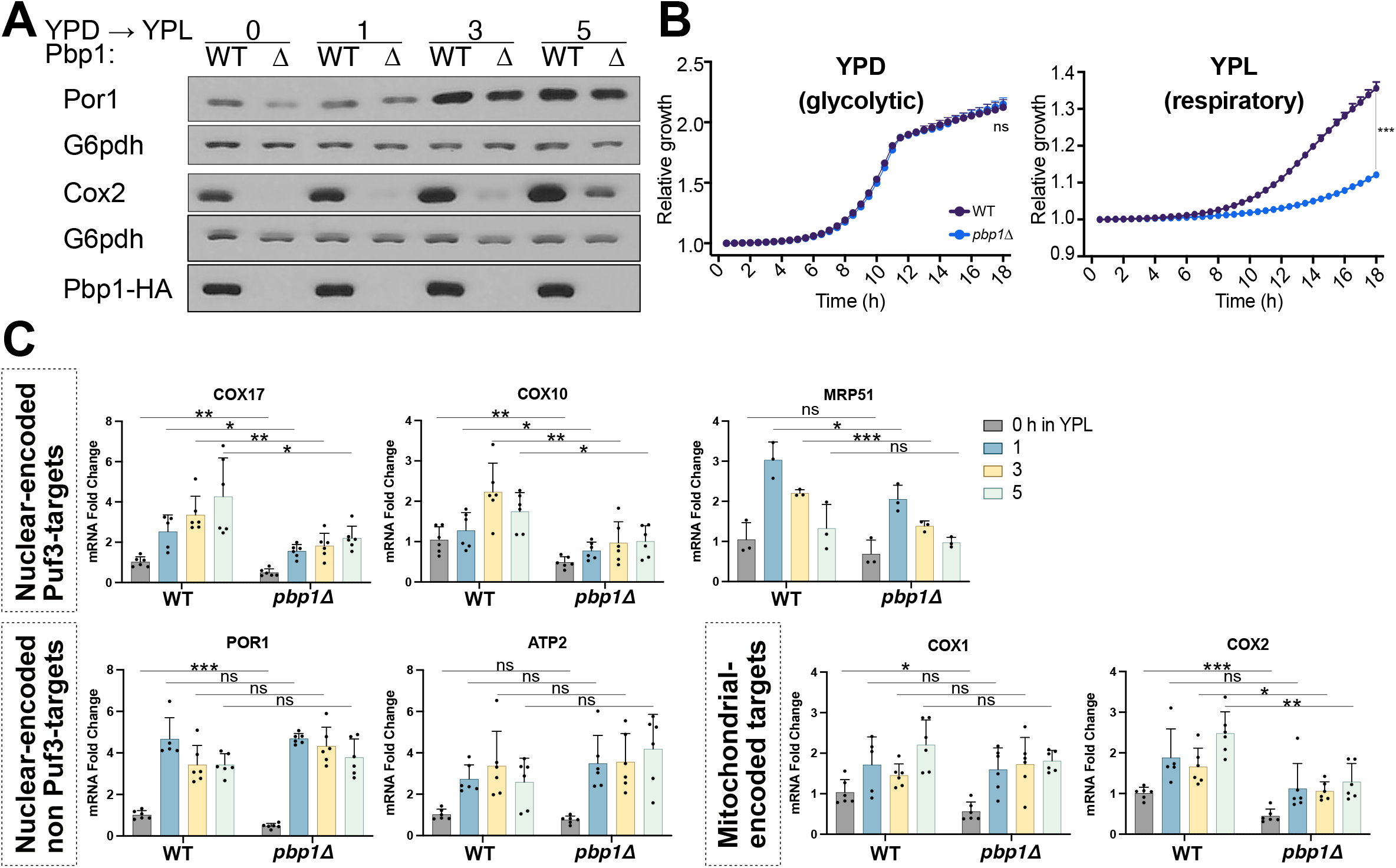
Pbp1 regulates Puf3-target mRNA and protein levels. A. A protein (Cox2) whose synthesis is dependent on Puf3-target mRNAs is decreased in *pbp1Δ* cells compared to WT. Cells were grown to log phase in glucose (YPD) media and then washed and resuspended in lactate (YPL) media. Samples were collected before and after switch to YPL, quenched, followed by protein extraction. Equals amounts of total protein were assessed by immunoblot for Cox2, Por1, Pbp1, and G6pdh levels. B. Cells lacking Pbp1 have a growth defect in respiratory (YPL) but not in glycolytic (YPD) growth conditions compared to WT cells. Cells were grown to log phase in glucose or lactate media, then resuspended at low OD in fresh media and A_600_ was measured every 30 min for 18 h. Error bars represent SD; n=3 and a two-tailed two-sample t-test, assuming equal variance was performed to determine statistical significance, p>0.05 (n.s.), ***p<0.001. C. Decreased mRNA levels of Puf3-target mRNAs are observed in *pbp1Δ* cells. The abundance of the indicated transcripts was measured by qRT-PCR and normalized to ACT1 levels at different time points after switching from YPD to YPL media. Error bars represent SD; n=6 except for MRP51; n=3. A two-tailed two-sample t-test, assuming equal variance, was used to determine statistical significance. p>0.05 (n.s.), *p<0.05, **p<0.01, ***p<0.001.

As Pbp1 interacts with Pab1 (poly(A)-binding protein 1) and contains putative RNA-binding domains (10), we tested the possibility that the loss of Pbp1 might affect the abundance of mRNAs involved in mitochondrial function. Analysis of a panel of nuclear- and mitochondrial-encoded mRNAs revealed that COX17, COX10, and MRP51 transcripts were decreased in *pbp1Δ* cells (Figure 1C). By contrast, no significant differences were observed in POR1 and ATP2 transcripts. Notably, the transcripts exhibiting reduced abundance in *pbp1Δ* mutant cells (COX17, COX10, MRP51) all have Puf3-binding elements in their 3’ UTRs and are Puf3-target mRNAs (6,9). For mitochondrial-encoded transcripts, we observed a modest decrease in COX2 transcripts in *pbp1Δ* cells at later time points in YPL and slight differences in COX1. The translation of COX2 relies on the mitochondrial ribosome, most subunits of which are post-transcriptional mRNA targets of Puf3 (e.g., MRP51). Therefore, defects in the translation of mitochondrial-encoded genes, such as Cox2, indicate disruption of Puf3 function (9).

The specific decrease in mRNA levels of Puf3-target mRNAs in *pbp1Δ* cells suggests that Pbp1 may help Puf3 stabilize and promote the translation of its target mRNAs. To test this possibility, we examined the extent of translation of a Puf3-binding element-containing mRNA following the shutoff of its transcription. The reporter consists of a mitochondrial targeting sequence (MTS), the GFP coding sequence, and the 3’ UTR of the mitochondrial ribosomal gene MRP51, which contains a Puf3-binding element (P3E) (Figure 2A) (9). The construct was integrated into a strain expressing a chimeric GAL4_DBD_-ER-VP16 transcription factor enabling inducible expression by the addition of ß-estradiol (16). This reporter allows monitoring of GFP transcript and protein amounts following a pulse of expression in the background of WT or *pbp1Δ* cells. Moreover, including a mutant reporter in which the P3E contains a four-nucleotide mutation allows the assessment of reporter translation dependent on an intact P3E.

**Figure 2.**
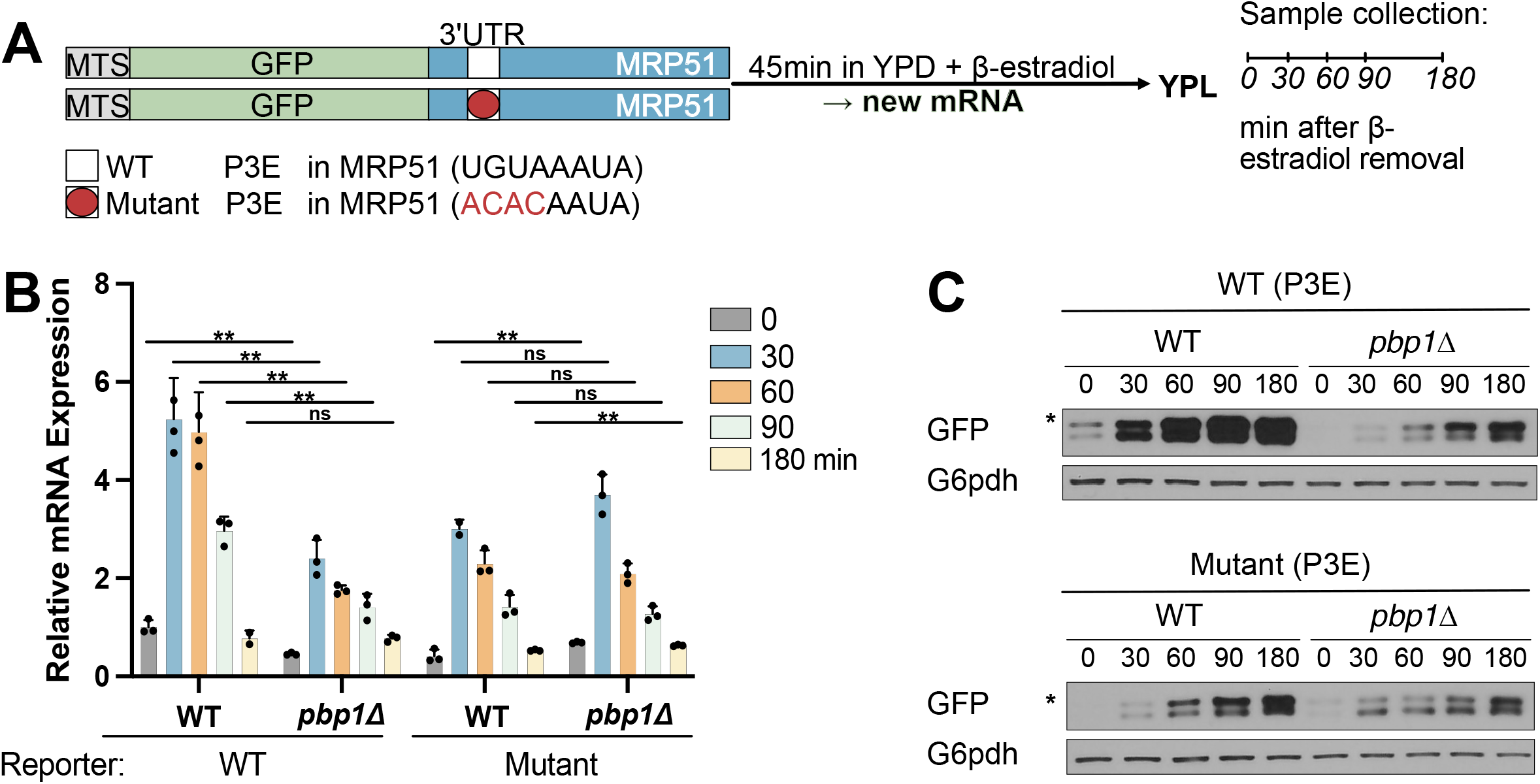
Pbp1 stabilizes and promotes the translation of Puf3-target mRNAs in respiratory conditions. A. Schematic representation of the Puf3 mRNA reporter assay. The indicated reporter constructs containing either a WT or mutant 3’ UTR of the MRP51 gene were placed under the control of the GAL1 promoter and integrated into strains expressing GAL4DBD-ER-VP16, in a WT or *pbp1Δ* background. After cells were grown in YPD to log phase, β-estradiol was added to induce a pulse of transcription of the reporter mRNA. After 45 min, cells were washed twice to remove β-estradiol and then resuspended in YPL medium. P3E = <u>P</u>uf<u>3</u>-binding <u>E</u>lement, MTS = mitochondrial targeting sequence. B. A reduction in GFP-mRNA can be observed for the reporter with intact P3E in *pbp1Δ* cells compared to WT, while no significant differences are observed for the mutant P3E reporter. Cells were harvested at 0, 30, 60, 90, and 180 min for analysis of reporter mRNA levels by qRT-PCR for GFP and normalized to ACT1. The increase in mRNA levels observed after medium switch is due to an unavoidable washing step to remove β-estradiol. Note that Puf3 is not absolutely required for the translation of its target mRNAs. Error bars represent SD; n=3 and a two-tailed two-sample t-test, assuming equal variance, was used to determine statistical significance. p>0.05 (n.s.), *p<0.05, **p<0.01. C. Pbp1 promotes the translation of the reporter containing an intact P3E. Reporter translation was assayed using an anti-GFP antibody, and G6pdh was used as a loading control. * denotes the slightly larger, unprocessed form of GFP containing a MTS.

WT and *pbp1Δ* cells were grown in YPD media, and ß-estradiol was added for 45 min before washout and then switched to YPL respiratory media. We observed the highest reporter mRNA expression at 30 min, followed by an expected, gradual decrease in mRNA due to the absence of any inducer (Figure 2B). Strikingly, levels of the reporter mRNA were significantly lower in *pbp1Δ* cells, strongly suggesting that Pbp1 is required to stabilize Puf3-target mRNAs. Interestingly, mRNA expression levels of the mutant P3E reporter were similar in WT and *pbp1Δ* cells, indicating that the reduced reporter expression in *pbp1Δ* cells is mediated by Puf3 and an intact P3E sequence in its target mRNAs.

Upon examination of translation of the reporter mRNA containing an intact P3E, WT cells synthesized significant amounts of GFP protein as a function of time in YPL media, despite decreasing reporter mRNA levels. In contrast, *pbp1*Δ mutant cells synthesized significantly lower amounts of GFP protein than WT (Figure 2C). No significant differences in translation of the reporter with a mutant P3E were observed. Interestingly, a basal amount of reporter translation was still observed in *pbp1Δ* cells. It has been suggested that Tom20p, a component of the outer mitochondrial membrane translocase complex, can recruit Puf3-target mRNAs to the mitochondrial surface even in the absence of Puf3 (17). Taken together, these data demonstrate that Pbp1 stabilizes Puf3-target mRNAs and promotes their translation into protein.

We next tested whether there might be a genetic interaction between Pbp1 and Puf3. We generated *puf3Δ, pbp1Δ*, and *pbp1Δpuf3Δ* cells and examined different mitochondrial protein levels by immunoblot following growth in YPD and YPL media (Figure 3A). As expected, Cox2, Por1, and Atp2 (subunit of the mitochondrial ATP synthase) protein levels all increased upon switching to respiratory conditions. However, Cox2 protein levels, which are dependent on the translation of Puf3-target mRNAs, decreased in *pbp1Δ, puf3Δ*, and *pbp1Δpuf3Δ* cells compared to WT cells. Por1 and Atp2 protein levels, which do not depend on Puf3-targets, were similar amongst the tested strains. Interestingly, the double knockout *pbp1Δpuf3Δ* exhibited a similar reduction in Cox2 as *puf3Δ* cells, suggesting that Puf3 is epistatic to Pbp1. In addition, we confirmed that Pbp1 levels were not affected by the absence of Puf3, whereas Puf3 levels were only modestly decreased in the absence of Pbp1 but only in lactate media (Figure S1). Taken together, these data suggest that Pbp1 and Puf3 function together in the same pathway and are consistent with the hypothesis that Pbp1 regulates Puf3. Notably, a decrease in each of the surveyed mitochondrial protein levels can also be observed in *pbp1Δ* cells in YPD, which is suggestive of a general impairment in mitochondria even in glucose conditions. However, for this study, we focus on the role of Pbp1 in respiratory conditions due to the exacerbated phenotype of reduced Cox2 levels and Puf3-target mRNA destabilization, specifically observed in YPL media.

**Figure 3.**
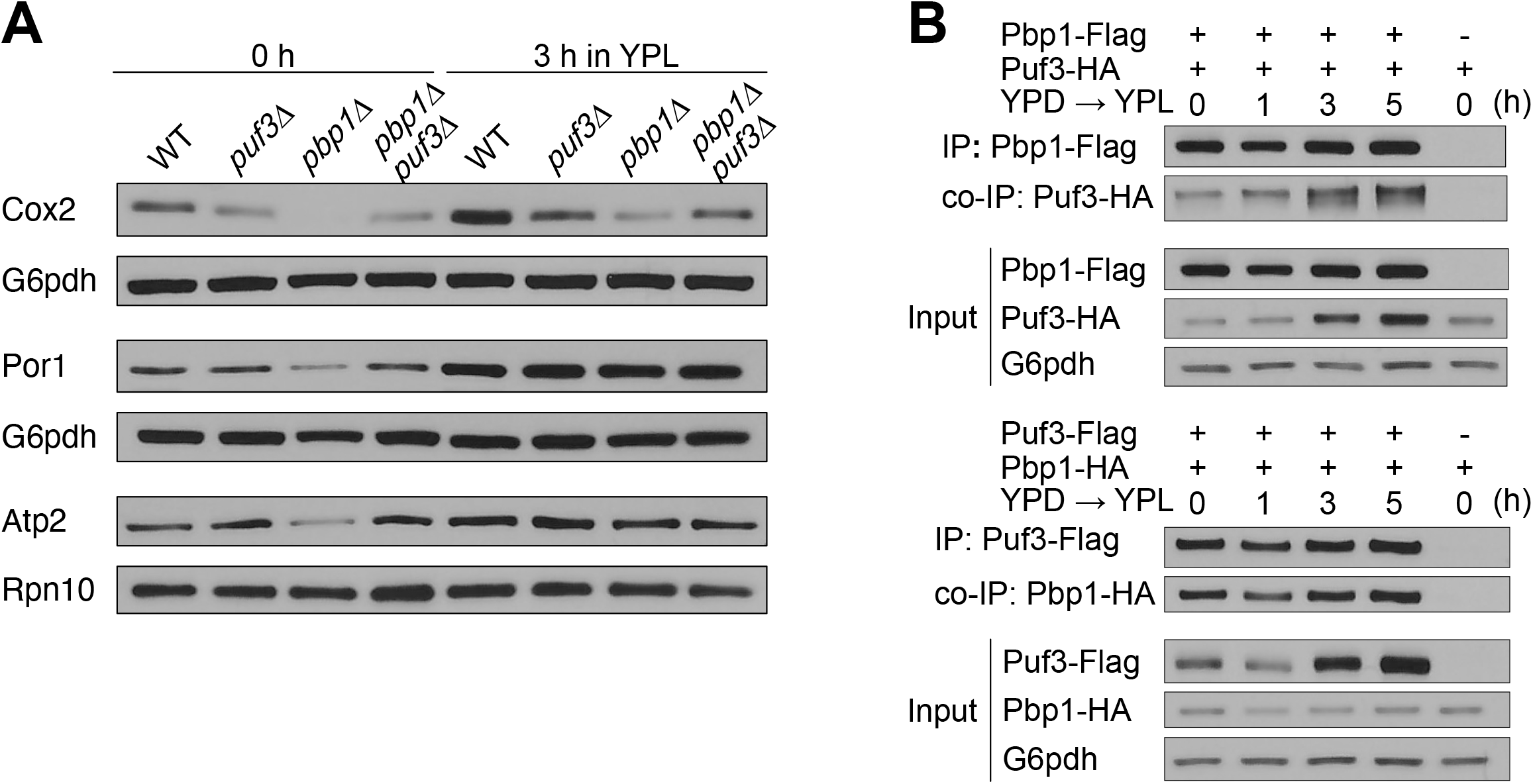
Increased amounts of Puf3 interact with Pbp1 under respiratory conditions. A. Only proteins whose synthesis is dependent on Puf3-target mRNAs (Cox2) are decreased in *puf3Δ, pbp1Δ*, and *pbp1Δpuf3Δ* cells as compared to WT. Cells were grown to log phase in YPD, then washed and resuspended in YPL. Samples were collected before (0 h) and after switch to YPL (3 h), quenched, followed by protein extraction. Equal amounts of protein were assayed by immunoblot for Cox2, Por1, Atp2, Rpn10 and G6pdh levels. B. Increased amounts of Puf3 are associated with Pbp1 in respiratory conditions. Cells with epitope-tagged Pbp1 and Puf3 were grown in YPD and then washed and resuspended in YPL. Samples were collected at the indicated time points, after which Flag immunoprecipitation was performed. – denotes negative control cells lacking a Flag-tag.

Having observed genetic interactions and phenotypic consequences on Puf3-targets in *pbp1Δ* mutant cells, we next investigated whether Pbp1 and Puf3 interact by co-immunoprecipitation analysis using C-terminal, epitope-tagged versions of Pbp1 (Pbp1-FLAG) and Puf3 (Puf3-HA) (Figure 3B). Pbp1-Flag co-IP experiments revealed that Pbp1 was indeed able to pull down Puf3. In addition, we performed the reciprocal Flag co-IP and obtained corresponding results, demonstrating that Pbp1 and Puf3 interact. We observed that Puf3 protein levels increase over time in response to the switch to respiratory conditions (Figure 3B and S1). Interestingly, we observed an increased interaction between Pbp1 and Puf3 over time in YPL media (Figure 3B). This suggests that the Pbp1-Puf3 interaction is responsive to the cellular metabolic state. These considerations led us to use respiratory growth conditions (3 h in YPL lactate media) in all subsequent experiments.

Previous studies have shown that Pbp1 is a negative regulator of TORC1 (14). We next tested whether the decreased Cox2 protein amounts in *pbp1*Δ cells might be due to increased TORC1 activity by treating cells with rapamycin. No difference in Cox2 levels was observed in *pbp1Δ* cells following rapamycin addition, indicating that the Cox2 protein reduction in *pbp1*Δ cells is independent of TORC1 signaling and instead likely due to its interaction with Puf3 (Figure S2).

To determine which region within Pbp1 is required for binding Puf3, we engineered a series of Pbp1 truncations (Figure 4A). Using co-immunoprecipitation, we observed that deletion of the mid-section (aa 299-570) or the low-complexity domain (aa 570-722) largely abolished the ability of Pbp1 to pull down Puf3 (Figure 4B). We then asked if the Pbp1-Puf3 interaction facilitates the translation of Puf3-target mRNAs by assessing Cox2 and Por1 protein levels in these mutants. We observed reduced Cox2, but not Por1, protein levels in Pbp1 *mid*Δ and Pbp1 *LCD*Δ cells, while no differences were observed in cells expressing the other Pbp1 truncation constructs (Figure 4C). Consistent with these effects on protein levels, qRT-PCR analysis showed reduced mRNA levels of Puf3-target transcripts COX17 and COX10, but not POR1 or ATP2, in the corresponding truncation mutants (Figure 4D).

**Figure 4.**
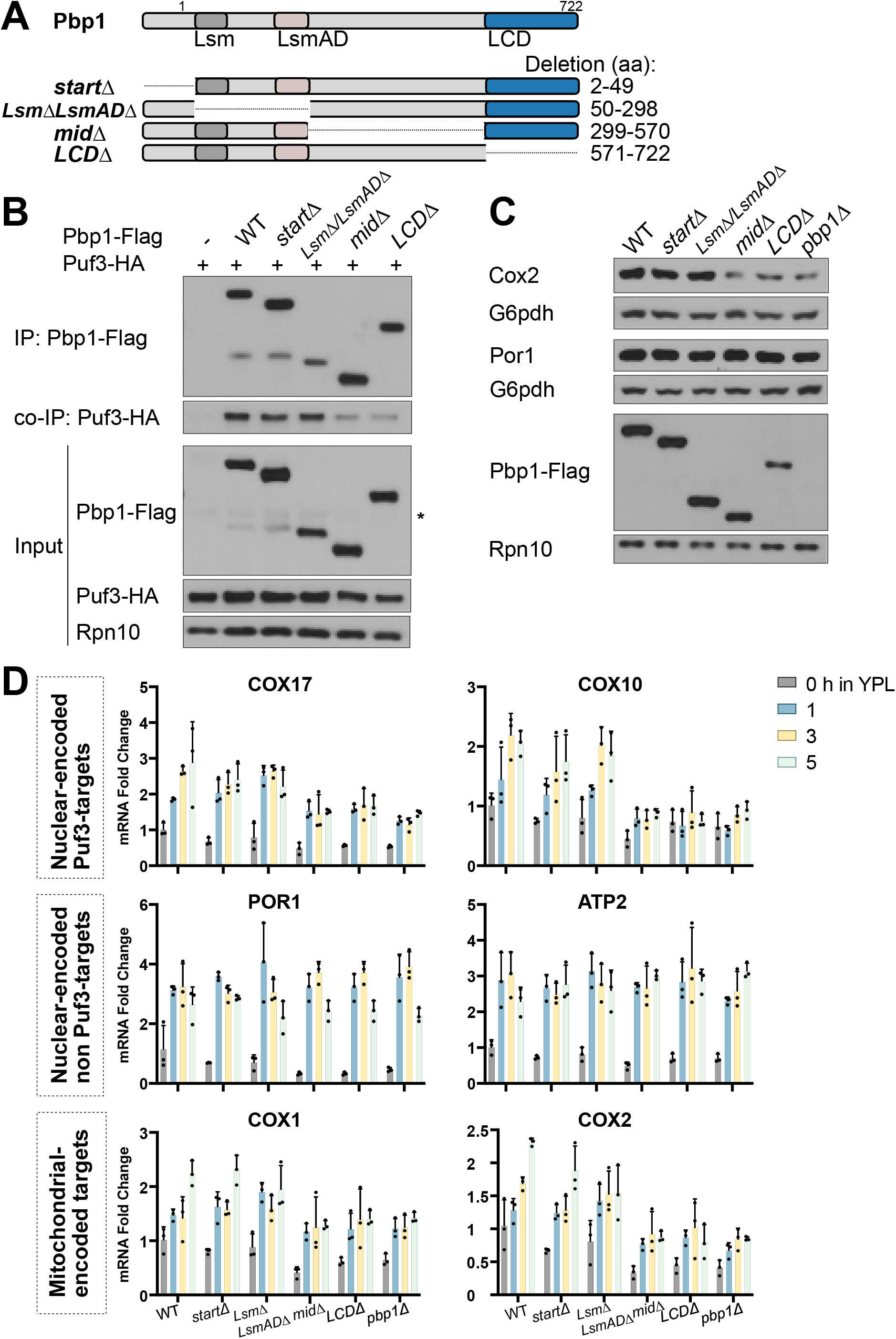
C-terminal, low complexity regions of Pbp1 mediate its interaction with Puf3. A. Schematic representation of Pbp1 and the deletion mutants used. B. Interaction between Puf3 and Pbp1 lacking its mid-section (*midΔ*) or C-terminal LCD (*LCDΔ*) is decreased in respiratory conditions. Cells with epitope-tagged Pbp1 and Puf3 were grown in YPD and then washed and resuspended in YPL. Samples were collected at the indicated time points, after which Flag immunoprecipitation was performed. – denotes negative control lacking a Flag-tag, but including HA-tag. * denotes non-specific band. C. Cox2 protein levels are decreased in Pbp1 *midΔ, LCDΔ*, and *pbp1Δ* in respiratory conditions. Cells were grown to log phase in YPD, then washed and resuspended in YPL. Samples were collected after 3 h in YPL, quenched, followed by protein extraction. Equal amounts of protein were assessed by immunoblot for Cox2, Por1, Atp2, Pbp1, Rpn10, and G6pdh levels. D. The abundance of the indicated transcripts was measured by qRT-PCR and normalized to ACT1 levels at different time points after switching from YPD to YPL. Error bars represent SD; n=3.

We previously found that Pbp1 self-associates through its low complexity domain, which is mediated by key methionine residues in the LCD (14,15). Replacement of eight methionine residues in the LCD by serine (M8S) leads to weaker self-assembly, increased TORC1 signaling, and reduced autophagy. In contrast, the M8F and M8Y mutants show stronger self-assembly and slightly increased autophagy (14,15). Therefore, we asked if these key methionine residues in the LCD responsible for the ability of Pbp1 to form condensates might also alter its interaction with Puf3 (Figures 5A and 5B). Interestingly, the M8S mutant showed reduced interaction with Puf3, in line with its compromised ability to phase separate and form assemblies. In contrast, the M8F and M8Y mutants showed increased interaction with Puf3, which is consistent with their ability to form stronger assemblies.

**Figure 5.**
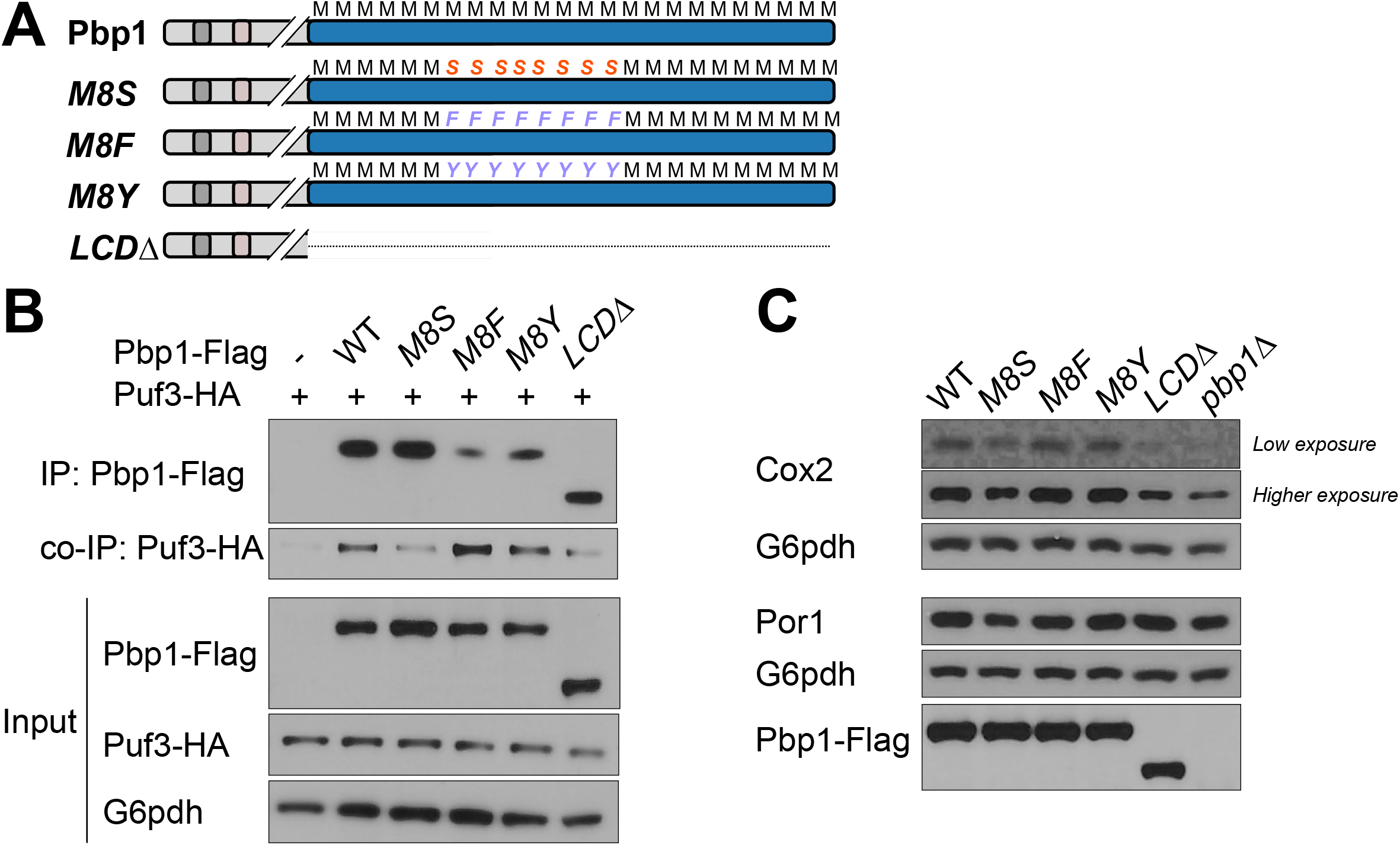
Methionine residues in the low complexity domain of Pbp1 are critical for its interaction with Puf3 and expression of Cox2 protein. A. Schematic representation of Pbp1 LCD-mutants used. Methionine residues within this key region of the LCD (M591, M595, M605, M606, M614, M616, M618, M625) were mutated to serine (M8S), phenylalanine (M8F), or tyrosine (M8Y). Pbp1 M8S forms weaker assemblies than WT, whereas M8F and M8Y both form stronger assemblies. B. Pbp1-Puf3 interaction is decreased in the M8S mutant. Cells expressing the indicated, epitope-tagged variants of Pbp1 and Puf3 were grown in YPD and then washed and resuspended in YPL. Samples were collected at 3 h in YPL, after which Flag-immunoprecipitation was performed. – denotes negative control lacking Flag-tag. C. Cox2 protein levels are decreased in M8S, LCD, and *pbp1Δ* mutants. Cells expressing the indicated epitope-tagged variants of Pbp1 were grown to log phase in YPD, then washed and resuspended in YPL. After 3 h, cells were harvested and quenched, followed by protein extraction. Equal amounts of protein were assessed by immunoblot for Cox2, Por1, Pbp1, and G6pdh levels.

Furthermore, Cox2 protein expression in these mutants, but not Por1, was affected in a manner that correlated with their ability to interact with Puf3. Decreased Cox2 protein levels were observed in the M8S mutant (Figure 5C), whereas increased Cox2 protein levels were observed in the M8F and M8Y mutants, compared to WT. These data suggest a strong correlation between the ability of Pbp1 to self-associate, the extent of its interaction with Puf3, and resulting amounts of Cox2 protein.

To further characterize the interaction between Pbp1 and Puf3, we used different truncation mutants of Puf3 (Figure 6A). Puf3 contains a PUF domain that binds to the 3’ UTR of mRNAs encoding mitochondrial proteins as well as a low-complexity N-terminal (Nt) region (7,9). It also contains a stretch of glutamine residues (polyQ) amid the protein. Puf3 *PUFΔ* and Puf3 *NtΔ* proteins were expressed at lower amounts than WT Puf3 or Puf3 *polyQ*Δ proteins (Figures 6B, C). Deleting the Nt region abolished the ability of Puf3 to pull down Pbp1. The Puf3 *PUFΔ* mutant also showed reduced interaction with Pbp1. In addition, Cox2 and Por1 protein levels were assayed in these mutants (Figure 6C). We observed reduced Cox2 but normal amounts of Por1 protein levels in Puf3 *NtΔ* and Puf3 *PUFΔ*, similar to *puf3Δ* cells, in YPL media. We then assayed mRNA levels by qRT-PCR analysis, and decreased amounts of Puf3-target mRNAs (e.g., COX10) were observed for the *NtΔ, PUFΔ*, and complete deletion (Figure S3). We conclude that without the N-terminal low complexity region or PUF domain, Puf3 can no longer interact with Pbp1 in respiratory conditions. Our cumulative findings suggest that Pbp1 and Puf3 interact via their low complexity domains (C-term LCD of Pbp1, N-term LCD of Puf3) to stabilize and promote translation of Puf3-target mRNAs for mitochondrial biogenesis.

**Figure 6.**
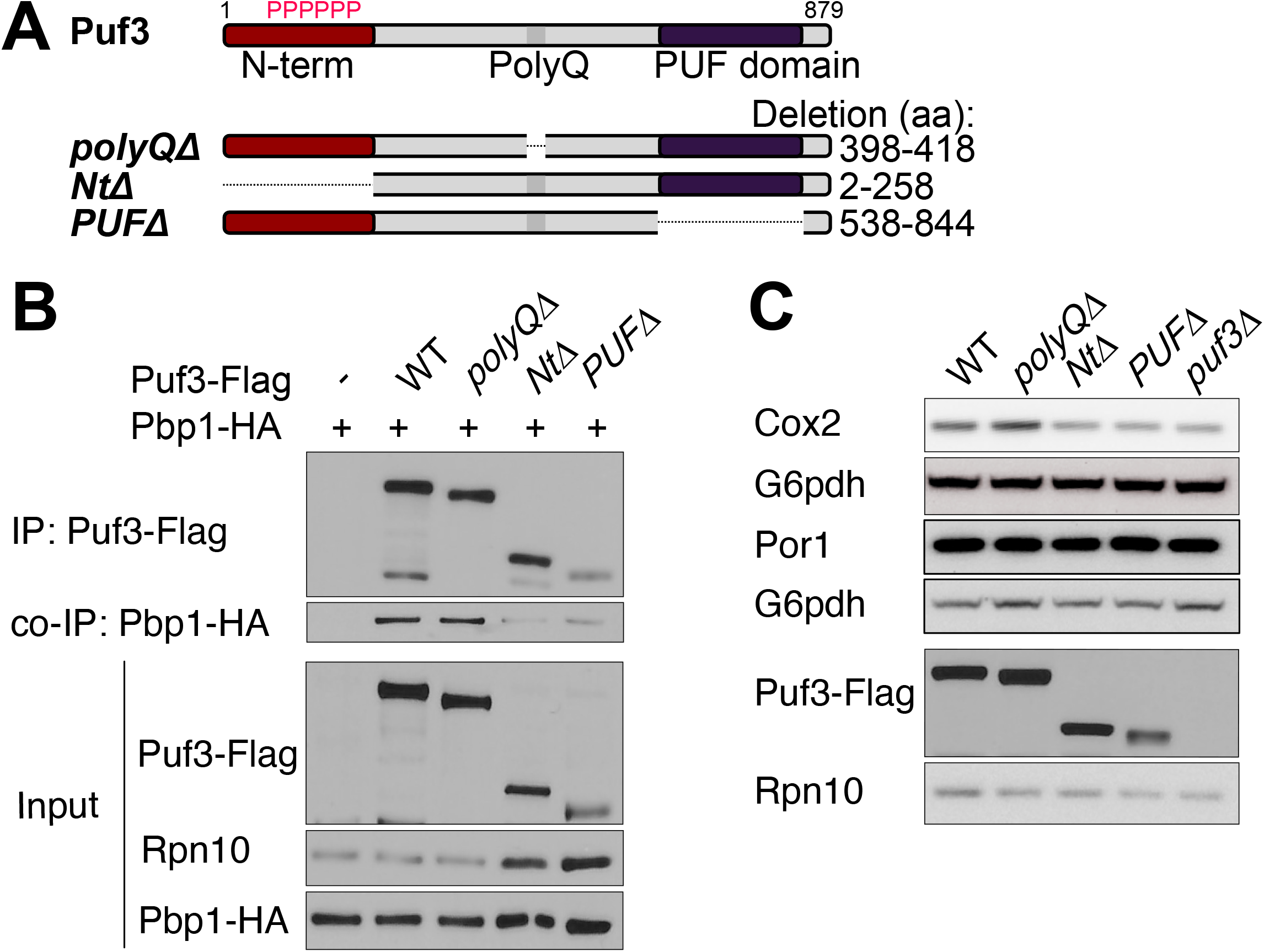
The N-terminal low complexity and PUF domains of Puf3 are critical for interaction with Pbp1. A. Schematic representation of Puf3 and deletion mutants used. B. Decreased interaction between Puf3 and Pbp1 is observed when the N-terminal LCD or PUF domain is lacking. Cells with endogenously tagged Pbp1 and Puf3 were grown in YPD and then washed and resuspended in YPL. Samples were collected after 3 h in YPL, after which Flag-immunoprecipitation was performed. – denotes negative control lacking Flag-tag. Note: total protein amounts are doubled for *Puf3 NtΔ* and *Puf3 PUFΔ* samples as compared to the others, as the expression of these two mutants is reduced (see Input). C. Cox2 protein levels are decreased in *Puf3 NtΔ* and *Puf3 PUFΔ* mutants in respiratory conditions. Cells were grown to log phase in YPD and then switched to YPL. Samples were collected after 3 h in YPL, quenched, followed by protein extraction. Equal amounts of protein were assessed by immunoblot for Cox2, Por1, Rpn10, and G6pdh levels. Note that *NtΔ* and *PUFΔ* mutants of Puf3 are unstable and expressed at lower levels compared to WT.

## Discussion

In this study, we show that Pbp1 supports mitochondrial function by promoting the stabilization and translation of Puf3-target mRNAs that are involved in mitochondrial biogenesis. Both Pbp1 and Puf3 harbor low complexity domains, and their association is required for normal Puf3 function. We speculate that Pbp1 self-associates under respiratory conditions to recruit Puf3 to the vicinity of mitochondria, where Puf3 promotes the translation of mRNAs central for mitochondrial biogenesis. Consistent with this hypothesis, we observed that Pbp1’s capacity to self-assemble correlated with its interaction with Puf3 and its ability to boost Cox2 protein amounts. Interestingly, Pbp1 has been suggested to function as a redox sensor (15). Reactive oxygen species can accumulate during mitochondrial dysfunction, underlining the potential for a key sensor role for Pbp1 in modulating Puf3 functions in response to mitochondrial dysfunction or biogenesis.

Pbp1 has been proposed to regulate mRNA decay and translation through interactions with various RNA regulatory factors including the poly(A) binding protein, the DEAD-box helicase Dhh1/DDX6, and the CCR4-Not and Pan2-Pan3 deadenylase complexes (10,18). This may represent a general mechanism to regulate mRNA decay. However, a recent study reported reduced COX10 and mitochondrial ribosomal protein mRNAs in the absence of Pbp1 in respiratory conditions, suggesting that Pbp1 exhibits specificity for particular transcripts (19). An increasing body of evidence links Pbp1 and mitochondrial respiratory functions, showing that cells without Pbp1 have increased petite formation and sensitivity to mitochondrial toxins (14,15). Additionally, Pbp1 overexpression can rescue mitochondrial stress in different settings (12,13). Our results support the idea that Pbp1 enables and facilitates mitochondrial biogenesis by stabilizing specifically Puf3-target mRNAs. Additional interacting partners of Pbp1 and Puf3 are likely involved in stabilizing and promoting the translation of these mRNAs. Further studies are needed to uncover other proteins that function together with Pbp1 and Puf3 to sustain mitochondrial functions.

Both Pbp1 and its mammalian ortholog ataxin-2 (ATXN2) harbor conserved domains (Lsm, LsmAD, a methionine-rich LCD), suggesting that both ataxin-2 and Pbp1 may play similar roles in the cell (20,21). Human sequencing studies have linked mutations in ATXN2 with neurodegenerative diseases, such as spinocerebellar ataxia type 2 and amyotrophic lateral sclerosis (22–24). Murine studies have shown that loss of ataxin-2 leads to reduced mitochondrial enzymes in the liver and downregulation of fatty acid metabolism pathways, citric acid cycle, and branched-chain amino acids metabolism in both the liver and cerebellum, suggesting that ataxin-2 regulates several metabolic pathways (25). How ataxin-2 affects different pathways could possibly be explained by the modulation of mRNA levels, such as those involved in the mitochondrial stress response and the mitochondrial quality control factor PTEN-induced kinase 1 (PINK1) (26). In addition, the C-terminal LCD of *Drosophila melanogaster* ataxin-2 was found to mediate RNP granule formation (27). As such, we speculate that ataxin-2, like Pbp1, might also regulate mRNAs targeted to the mitochondria through interactions with its C-terminal LCD.

Neurons have a high mitochondrial density and require the maintenance of a functional mitochondrial population throughout cell bodies (28). The presence of mitochondrial dysfunction in neurons is frequently observed in neurodegenerative diseases, either as a hallmark or a consequence of the disease (29). Our work uncovering the function of Pbp1 in mitochondrial biogenesis strongly supports the investigation of ataxin-2 as a regulator of mitochondrial functions in the context of neurodegenerative conditions.

## Methods

### Yeast strains, growth, and media

The prototrophic *Saccharomyces cerevisiae* CEN.PK strain (30) was used in all experiments. All strains used in this study are listed in Table S1. Gene deletions were performed using standard PCR-based strategies to amplify resistance cassettes with appropriate flanking sequences and replace the target gene through homologous recombination (31). C-terminal tags were similarly made using PCR to amplify resistance cassettes with flanking sequences. Pbp1 and Puf3 mutants with various domain deletions or point mutations were first made using PCR and then integrated into the PBP1 or PUF3 locus in a *pbp1Δ* or *puf3Δ* strain with different selection markers. Yeast strains were grown in YPD (1% yeast extract (Bio Basic), 2% peptone (BD Biosciences) and 2% glucose) or YPL (0.5% yeast extract, 2% peptone and 2% lactate (Sigma L1375)). Cells from overnight cultures were inoculated into fresh YPD to 0.3 optical density (OD_600_)/ml and grown for at least two generations to log phase. Cells were then spun down, washed with YPL, and resuspended in the same volume of YPL. Samples were collected at indicated time points. For cells treated with rapamycin, 200 ng/ml rapamycin (Sigma) was added to cells grown in YPL and incubated for 30 min before harvesting.

### Growth curves

Cells were inoculated in YPD and grown overnight. The following day cells were resuspended in fresh media, YPD or YPL, and grown for ∼6 h. Cells were then diluted to an OD_600_ of 0.05 in fresh YPD or 0.1 in fresh YPL, and 100 µl of culture was pipetted into a 96 multiwell flat-bottom transparent plate. The plate was shaken and kept at 30°C while the absorbance was measured every 30 minutes for 18 hours by the SPARK multimode plate reader (TECAN) according to the manufacturer’s instructions.

### Whole-cell lysate protein extraction

Cell pellets (5 units of OD_600_) were quenched in 15% TCA for 15 min on ice, pelleted, and washed with ice-cold 100% acetone. Pellets were then resuspended in 240µl urea extraction buffer (6 M urea, 50 mM Tris-HCl pH 6.8, 1 mM PMSF, 1% SDS, 1 mM EDTA, 1 mM DTT, 1 mM Na_3_VO_4_×2H_2_O, 5 µM pepstatin A, 10 µM leupeptin, 2X Roche protease inhibitor cocktail), 200µl glass beads were added, and cells were lysed by bead-beating. After a 5 min incubation at 65°C, the lysate was centrifuged at maximum speed for 10 min. The supernatant was collected, and the protein concentration was normalized using a BCA protein assay kit (Thermo Scientific). Sample buffer (50 mM Tris-HCl pH 6.8, 2% SDS, 10% glycerol, 10 mM DTT, bromophenol blue) was added.

### Co-immunoprecipitation

At the indicated time points, 50 units of OD_600_ cells were harvested, flash-frozen with liquid nitrogen, and stored at -80°C until cell lysis. The cell pellet was resuspended in 350 µL of lysis buffer A (50 mM HEPES pH 7.5, 150 mM NaCl, 10 mM MgCl_2_, 0.5% NP-40, 2X protease inhibitor cocktail (Roche), 1 mM PMSF, 1 mM Na_3_VO_4_×2H_2_O, 50 µM pepstatin A, 10 µM leupeptin, 5 mM NaF). After adding 300µL of glass beads, cells were lysed by bead beating six times: 30 sec of beating / 2 min of cooling on ice. The lysed cells were then separated from glass beads by centrifugation at 6000 rpm for 2 min at 4°C and diluted with 525 µL of lysis buffer B (buffer A devoid of NP-40). Two successive centrifugations then clarified crude cell extracts at a maximum speed of 10 min at 4°C. The protein concentration of the cleared lysates was then measured using Bradford assay (Bio-Rad) and adjusted to be equal among all samples in 800 µL reaction volume. For input samples, 10 µL of the reactions was mixed with 30 µl 8 M urea and 12.5 µl 4X SDS sample dye with 40 mM DTT, denatured for 5 min at 65°C. For each co-immunoprecipitation reaction, 25 µL of Dynabeads Protein G (Life technologies) was washed with IP-lysis buffer and incubated with 3 µg of mouse anti-Flag antibody (Sigma, M2) for 1 h at 4°C. Unbound antibody was removed by centrifugation at 500 g for 1 min at 4°C. The conjugated dynabeads-antibody were then added to the cleared lysates. After incubating for 1-2 h at 4°C, the dynabeads were washed three times with wash buffer (50 mM HEPES pH 7.5, 150 mM NaCl, 1 mM EDTA, 2X protease inhibitor cocktail, 0.2% NP-40). The sample was collected by boiling the beads in 1X SDS sample dye with 6% β-mercaptoethanol for 5 min at 95°C.

### Western blot

Western blots were carried out by running whole cell lysate or co-IP samples on a NuPAGE 4-12% BisTris gel (ThermoFisher) with MOPS running buffer. Wet transfers at 350 mA at 4°C for 75-90 min were used to transfer protein onto a 0.45-micron nitrocellulose membrane. Membranes were blocked with 5% milk in TBST for 1 h and incubated with primary antibody overnight at 4°C in 1% milk in TBST. Membranes were then washed with TBST and incubated for 1 h with anti-mouse or anti-rabbit HRP antibody in 1% milk in TBST before incubating with Pierce ECL Western blotting substrate and exposing to film. The following antibodies were used: rabbit anti-Flag M2 (Cell Signaling, #2368), rabbit anti-Ha (Cell Signaling, #3724), rabbit anti-G6pdh ab2 (Sigma, #A9521), rabbit anti-Rpn10 (Abcam, #ab98843), mouse anti-Cox2 (ThermoFisher, #459150), mouse anti-Por1 (ThermoFisher, #459500), mouse anti-GFP (Roche, clone 7.1 and 13.1) and rabbit anti-Atp2 (32) (this antibody was a kind gift from Dr. S.M. Claypool at Johns Hopkins University).

### qRT-PCR analysis

Approximately 1 OD_600_ of cells was collected, flash-frozen, and stored at -80°C at the indicated time points. Total RNA was extracted with MasterPure Yeast RNA Purification Kit following the manufacturer’s protocol. cDNA was made with Superscript III reverse transcriptase (Invitrogen), and gene expression levels were analyzed by qRT-PCR with SYBR Green (Invitrogen). mRNA levels were normalized against ACT1. Primers are listed in Table S2.

### β-estradiol inducible reporter system

Experiments with the Puf3-regulated mRNA reporter were performed as previously described (9). In brief, cells were grown in YPD to log phase and 100 nM of β-estradiol was added for 45 min to induce transcription. Cells were then spun down and washed twice with YPD or YPL to remove residual β-estradiol. Cells were then resuspended in similar volume in fresh YPD or YPL and collected at 0, 30, 60, 90, and 180 min to assess the yEGFP mRNA and protein levels.

**Table 1:**
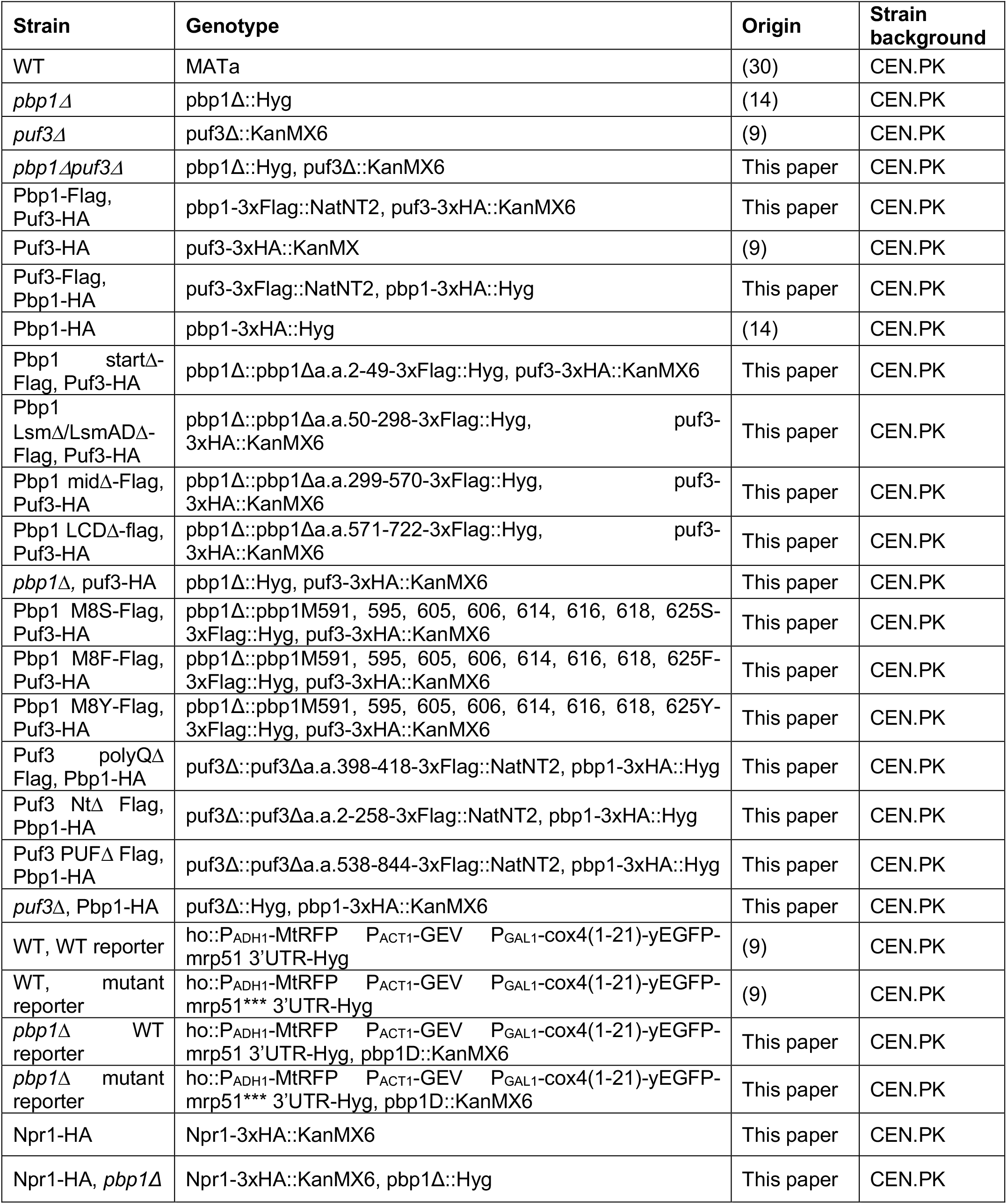
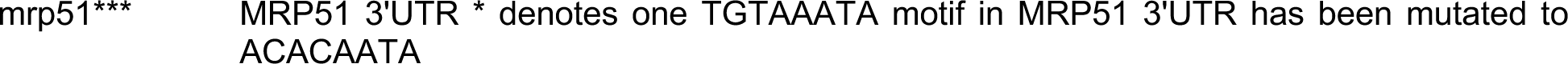
*Saccharomyces cerevisiae* strains used in this study.

| Strain | Genotype | Origin | Strain background |
| --- | --- | --- | --- |
| WT | MATa | (30) | CEN.PK |
| *pbp1Δ* | *pbp1Δ::Hyg* | (14) | CEN.PK |
| *puf3Δ* | *puf3Δ::KanMX6* | (9) | CEN.PK |
| *pbp1Δpuf3Δ* | *pbp1Δ::Hyg, puf3Δ::KanMX6* | This paper | CEN.PK |
| Pbp1-Flag, Puf3-HA | *pbp1-3xFlag::NatNT2, puf3-3xHA::KanMX6* | This paper | CEN.PK |
| Puf3-HA | *puf3-3xHA::KanMX* | (9) | CEN.PK |
| Puf3-Flag, Pbp1-HA | *puf3-3xFlag::NatNT2, pbp1-3xHA::Hyg* | This paper | CEN.PK |
| Pbp1-HA | *pbp1-3xHA::Hyg* | (14) | CEN.PK |
| Pbp1<sup>startΔ</sup>-Flag, Puf3-HA | *pbp1Δ::pbp1Δa.a.2-49-3xFlag::Hyg, puf3-3xHA::KanMX6* | This paper | CEN.PK |
| Pbp1 LsmΔ/LsmADΔ-Flag, Puf3-HA | *pbp1Δ::pbp1Δa.a.50-298-3xFlag::Hyg, puf3-3xHA::KanMX6* | This paper | CEN.PK |
| Pbp1 midΔ-Flag, Puf3-HA | *pbp1Δ::pbp1Δa.a.299-570-3xFlag::Hyg, puf3-3xHA::KanMX6* | This paper | CEN.PK |
| Pbp1 LCDΔ-flag, Puf3-HA | *pbp1Δ::pbp1Δa.a.571-722-3xFlag::Hyg, puf3-3xHA::KanMX6* | This paper | CEN.PK |
| *pbp1Δ, puf3-HA* | *pbp1Δ::Hyg, puf3-3xHA::KanMX6* | This paper | CEN.PK |
| Pbp1 M8S-Flag, Puf3-HA | *pbp1Δ::pbp1M591, 595, 605, 606, 614, 616, 618, 625S-3xFlag::Hyg, puf3-3xHA::KanMX6* | This paper | CEN.PK |
| Pbp1 M8F-Flag, Puf3-HA | *pbp1Δ::pbp1M591, 595, 605, 606, 614, 616, 618, 625F-3xFlag::Hyg, puf3-3xHA::KanMX6* | This paper | CEN.PK |
| Pbp1 M8Y-Flag, Puf3-HA | *pbp1Δ::pbp1M591, 595, 605, 606, 614, 616, 618, 625Y-3xFlag::Hyg, puf3-3xHA::KanMX6* | This paper | CEN.PK |
| Puf3 polyQΔ Flag, Pbp1-HA | *puf3Δ::puf3Δa.a.398-418-3xFlag::NatNT2, pbp1-3xHA::Hyg* | This paper | CEN.PK |
| Puf3 NtΔ Flag, Pbp1-HA | *puf3Δ::puf3Δa.a.2-258-3xFlag::NatNT2, pbp1-3xHA::Hyg* | This paper | CEN.PK |
| Puf3 PUFΔ Flag, Pbp1-HA | *puf3Δ::puf3Δa.a.538-844-3xFlag::NatNT2, pbp1-3xHA::Hyg* | This paper | CEN.PK |
| *puf3Δ, Pbp1-HA* | *puf3Δ::Hyg, pbp1-3xHA::KanMX6* | This paper | CEN.PK |
| WT, WT reporter | *ho::P<sub>ADH1</sub>-MtRFP P<sub>ACT1</sub>-GEV P<sub>GAL1</sub>-cox4(1-21)-yEGFP-mrp51 3'UTR-Hyg* | (9) | CEN.PK |
| WT, mutant reporter | *ho::P<sub>ADH1</sub>-MtRFP P<sub>ACT1</sub>-GEV P<sub>GAL1</sub>-cox4(1-21)-yEGFP-mrp51\*\*\* 3'UTR-Hyg* | (9) | CEN.PK |
| *pbp1Δ* WT reporter | *ho::P<sub>ADH1</sub>-MtRFP P<sub>ACT1</sub>-GEV P<sub>GAL1</sub>-cox4(1-21)-yEGFP-mrp51 3'UTR-Hyg, *pbp1D::KanMX6** | This paper | CEN.PK |
| *pbp1Δ* mutant reporter | *ho::P<sub>ADH1</sub>-MtRFP P<sub>ACT1</sub>-GEV P<sub>GAL1</sub>-cox4(1-21)-yEGFP-mrp51\*\*\* 3'UTR-Hyg, *pbp1D::KanMX6** | This paper | CEN.PK |
| Npr1-HA | *Npr1-3xHA::KanMX6* | This paper | CEN.PK |
| Npr1-HA, *pbp1Δ* | *Npr1-3xHA::KanMX6, pbp1Δ::Hyg* | This paper | CEN.PK |

mrp51\*\*\* MRP51 3'UTR \* denotes one TGTAATA motif in MRP51 3'UTR has been mutated to ACACAATA

**Table 2:**
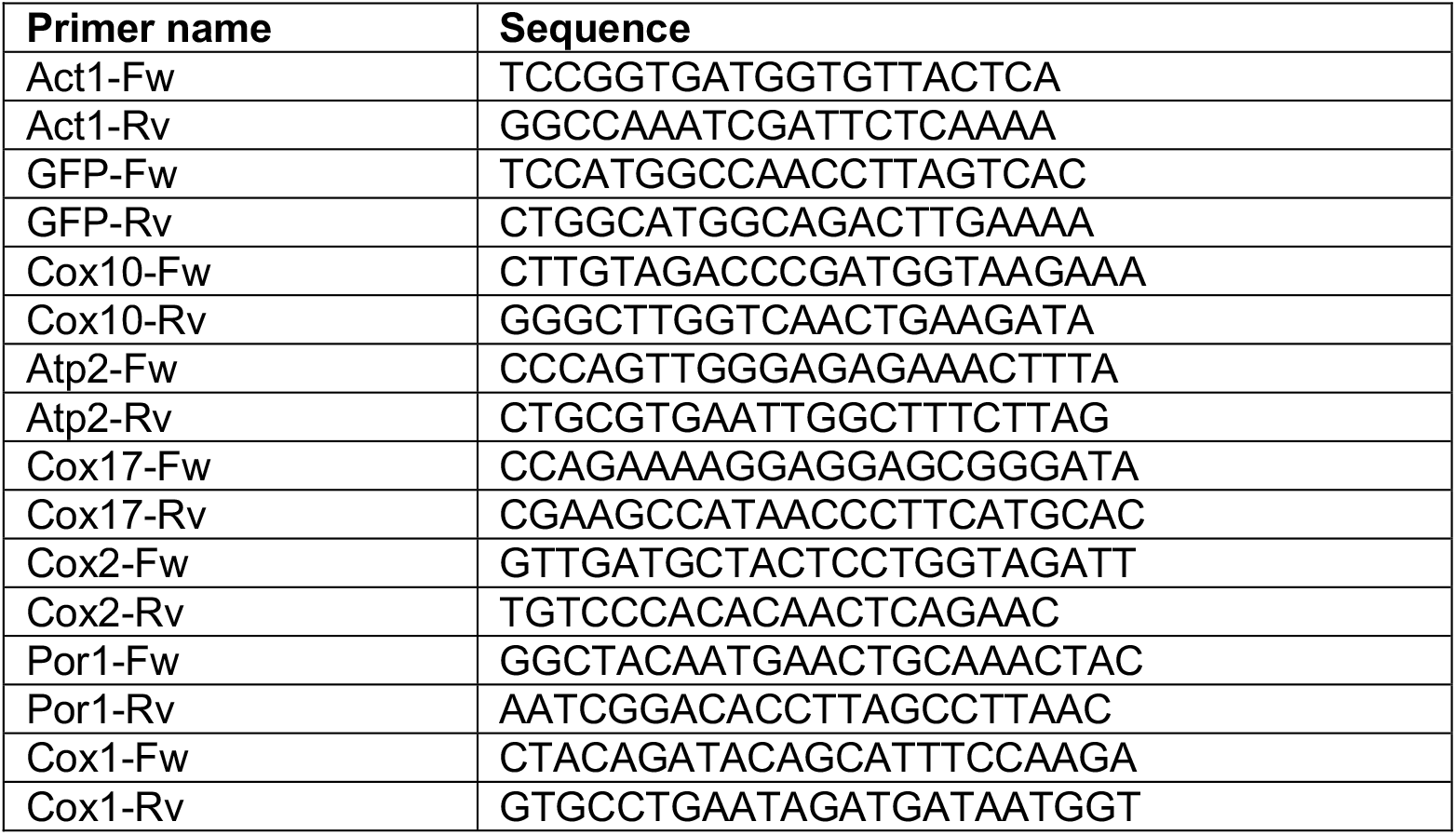
qRT-PCR primers.

## Acknowledgments

We thank the Tu Lab for helpful discussions, and D. Caballero, M.G. Acoba for helpful edits to the text. We thank Dr. S.M. Claypool at Johns Hopkins University for the Atp2 antibody. This work was supported by NIH grant R01NS115546. B.P.T. is a Howard Hughes Medical Institute Investigator at UT Southwestern.

## Supplemental Data

Supplemental Figures 1, 2 and 3.

**Supplemental Figure 1.**
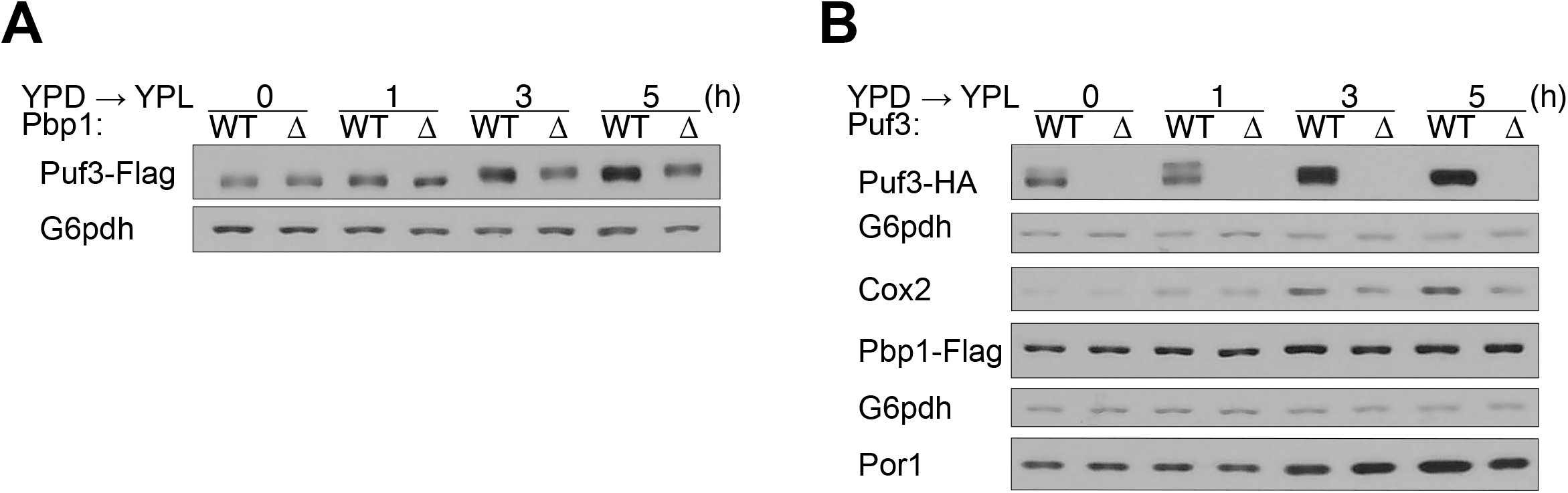
A. Puf3 protein levels are decreased in cells lacking Pbp1. Cells expressing Puf3-Flag were assayed in the presence or absence of Pbp1 for the indicated proteins by Western blot. B. Pbp1 protein levels are not affected in cells lacking Puf3. Cells expressing Pbp1-Flag were assayed in the presence or absence of Puf3 for the indicated proteins by Western blot at various time points following switch from YPD to YPL.

**Supplemental Figure 2.**
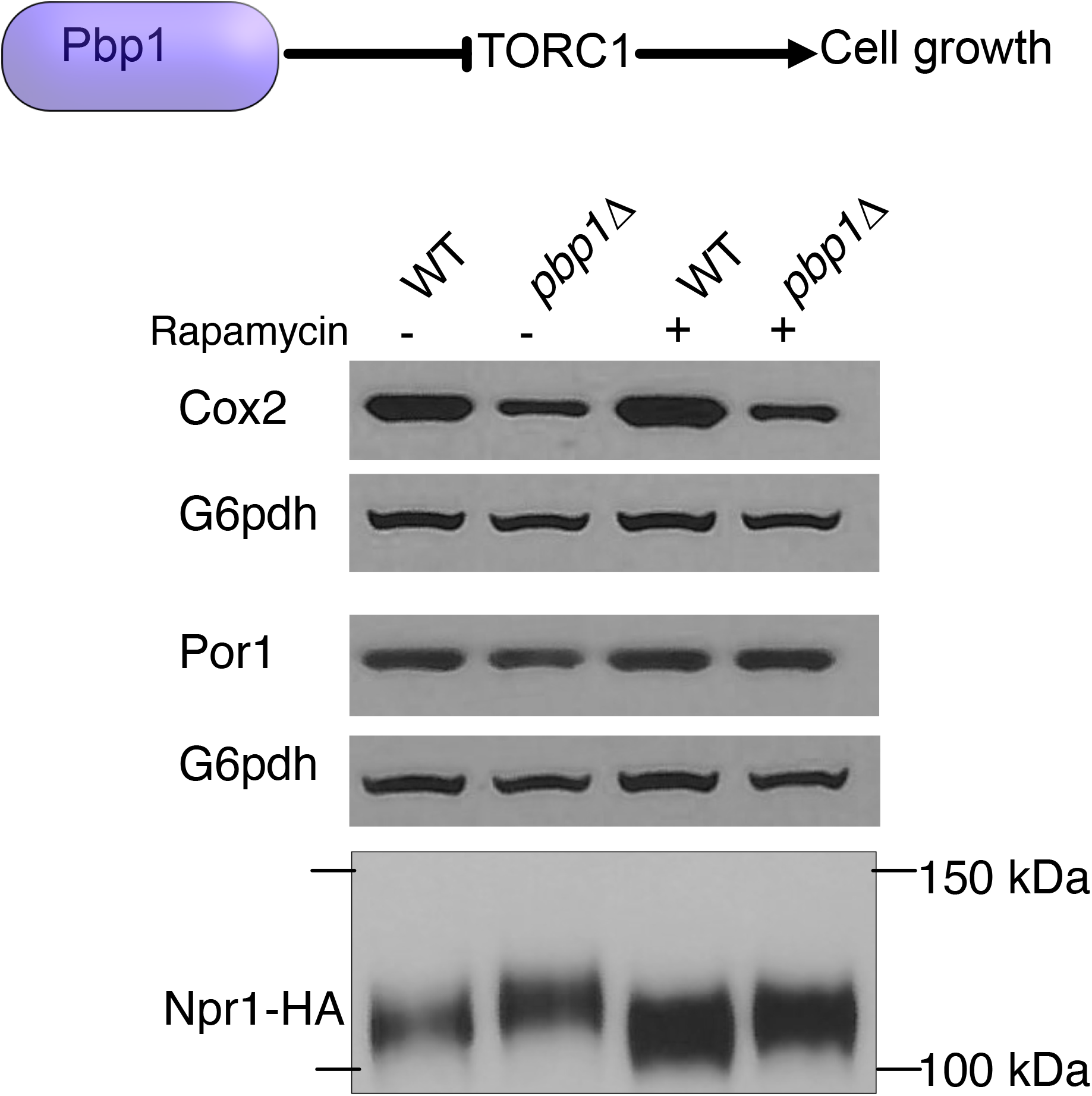
Decreased Cox2 levels in *pbp1Δ* cells are not merely due to hyperactive TORC1 signaling. The indicated strains were grown in YPL for 3 h, of which the last 30 min were in the absence (-) or presence (+) of 200 ng/mL rapamycin. Samples were collected, quenched, and extracted for immunoblot analysis of the indicated proteins. Note that rapamycin treatment did not restore protein amounts of Cox2 in *pbp1Δ* cells.

**Supplemental Figure 3.**
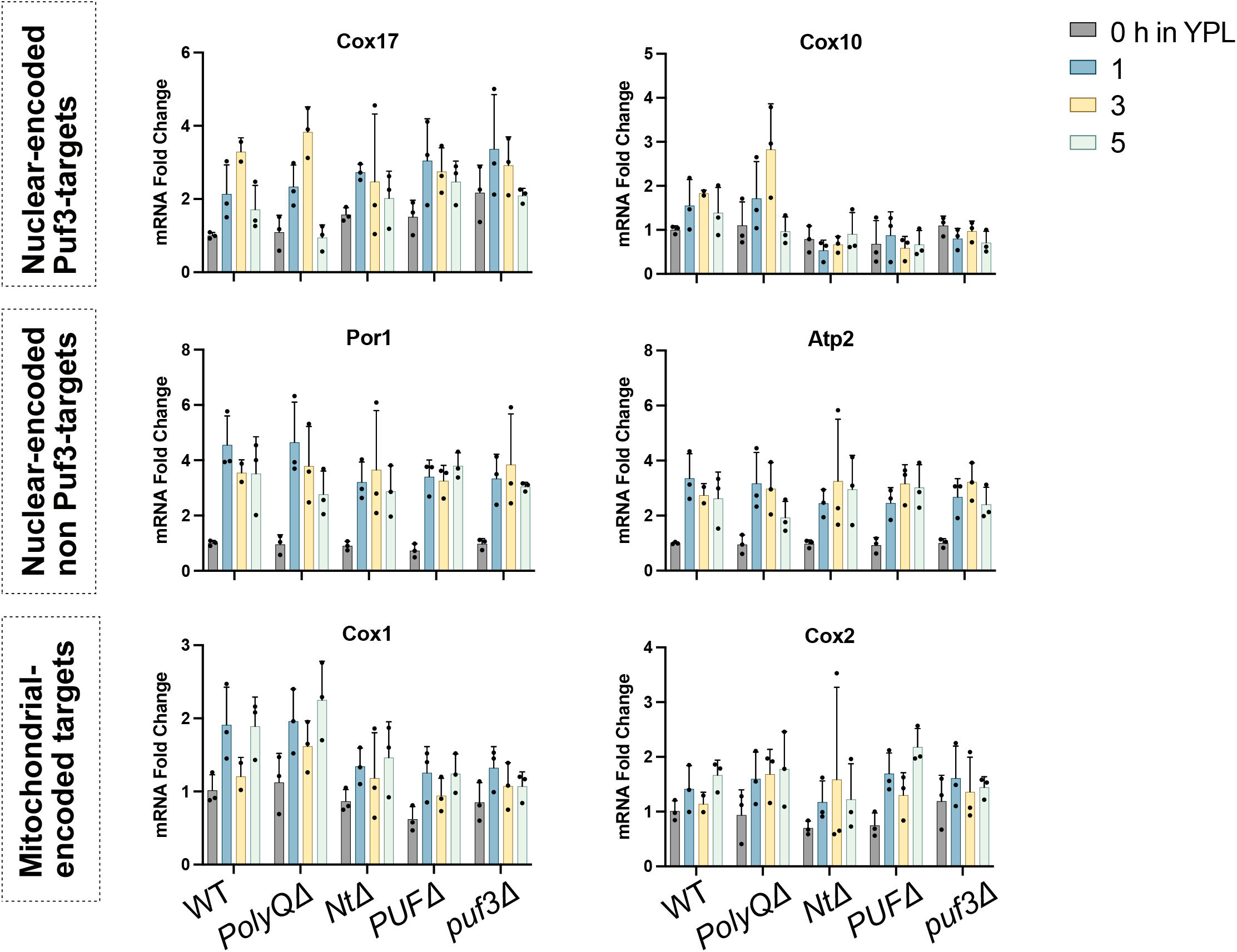
Several mRNAs involved in mitochondrial function were assayed by qRT-PCR in the indicated Puf3 mutant strains at the indicated time points after switch from YPD to YPL. The abundance of the indicated transcripts was normalized to ACT1.

## References

1. Couvillion MT, Soto IC, Shipkovenska G, Churchman LS. Synchronized mitochondrial and cytosolic translation programs. Nature. 2016 May 26;533(7604):499–503.

2. Nishanth MJ, Simon B. Functions, mechanisms and regulation of Pumilio/Puf family RNA binding proteins: a comprehensive review. Mol Biol Rep. 2020 Jan;47(1):785–807.

3. Chatenay-Lapointe M, Shadel GS. Repression of mitochondrial translation, respiration and a metabolic cycle-regulated gene, SLF1, by the yeast Pumilio-family protein Puf3p. PLoS One. 2011;6(5):e20441.

4. García-Rodríguez LJ, Gay AC, Pon LA. Puf3p, a Pumilio family RNA binding protein, localizes to mitochondria and regulates mitochondrial biogenesis and motility in budding yeast. J Cell Biol. 2007 Jan 15;176(2):197–207.

5. Gerber AP, Herschlag D, Brown PO. Extensive association of functionally and cytotopically related mRNAs with Puf family RNA-binding proteins in yeast. PLoS Biol. 2004 Mar;2(3):E79.

6. Lapointe CP, Stefely JA, Jochem A, Hutchins PD, Wilson GM, Kwiecien NW, et al. Multi-omics Reveal Specific Targets of the RNA-Binding Protein Puf3p and Its Orchestration of Mitochondrial Biogenesis. Cell Syst. 2018 Jan 24;6(1):125-135.e6.

7. Olivas W, Parker R. The Puf3 protein is a transcript-specific regulator of mRNA degradation in yeast. EMBO J. 2000 Dec 1;19(23):6602–11.

8. Tu BP, Kudlicki A, Rowicka M, McKnight SL. Logic of the yeast metabolic cycle: temporal compartmentalization of cellular processes. Science. 2005 Nov 18;310(5751):1152–8.

9. Lee CD, Tu BP. Glucose-Regulated Phosphorylation of the PUF Protein Puf3 Regulates the Translational Fate of Its Bound mRNAs and Association with RNA Granules. Cell Rep. 2015 Jun 16;11(10):1638–50.

10. Mangus DA, Amrani N, Jacobson A. Pbp1p, a factor interacting with Saccharomyces cerevisiae poly(A)-binding protein, regulates polyadenylation. Mol Cell Biol. 1998 Dec;18(12):7383–96.

11. Swisher KD, Parker R. Localization to, and effects of Pbp1, Pbp4, Lsm12, Dhh1, and Pab1 on stress granules in Saccharomyces cerevisiae. PLoS One. 2010 Apr 2;5(4):e10006.

12. Dunn CD, Jensen RE. Suppression of a defect in mitochondrial protein import identifies cytosolic proteins required for viability of yeast cells lacking mitochondrial DNA. Genetics. 2003 Sep;165(1):35–45.

13. Wang X, Chen XJ. A cytosolic network suppressing mitochondria-mediated proteostatic stress and cell death. Nature. 2015 Aug 27;524(7566):481–4.

14. Yang YS, Kato M, Wu X, Litsios A, Sutter BM, Wang Y, et al. Yeast Ataxin-2 Forms an Intracellular Condensate Required for the Inhibition of TORC1 Signaling during Respiratory Growth. Cell. 2019 Apr 18;177(3):697-710.e17.

15. Kato M, Yang YS, Sutter BM, Wang Y, McKnight SL, Tu BP. Redox State Controls Phase Separation of the Yeast Ataxin-2 Protein via Reversible Oxidation of Its Methionine-Rich Low-Complexity Domain. Cell. 2019 Apr 18;177(3):711-721.e8.

16. McIsaac RS, Silverman SJ, McClean MN, Gibney PA, Macinskas J, Hickman MJ, et al. Fast-acting and nearly gratuitous induction of gene expression and protein depletion in Saccharomyces cerevisiae. Mol Biol Cell. 2011 Nov;22(22):4447–59.

17. Eliyahu E, Pnueli L, Melamed D, Scherrer T, Gerber AP, Pines O, et al. Tom20 mediates localization of mRNAs to mitochondria in a translation-dependent manner. Mol Cell Biol. 2010 Jan;30(1):284–94.

18. Mangus DA, Smith MM, McSweeney JM, Jacobson A. Identification of factors regulating poly(A) tail synthesis and maturation. Mol Cell Biol. 2004 May;24(10):4196–206.

19. Tuong Vi DT, Fujii S, Valderrama AL, Ito A, Matsuura E, Nishihata A, et al. Pbp1, the yeast ortholog of human Ataxin-2, functions in the cell growth on non-fermentable carbon sources. PLoS One. 2021;16(5):e0251456.

20. Albrecht M, Golatta M, Wüllner U, Lengauer T. Structural and functional analysis of ataxin-2 and ataxin-3. Eur J Biochem. 2004 Aug;271(15):3155–70.

21. Albrecht M, Lengauer T. Survey on the PABC recognition motif PAM2. Biochem Biophys Res Commun. 2004 Mar 26;316(1):129–38.

22. Imbert G, Saudou F, Yvert G, Devys D, Trottier Y, Garnier JM, et al. Cloning of the gene for spinocerebellar ataxia 2 reveals a locus with high sensitivity to expanded CAG/glutamine repeats. Nat Genet. 1996 Nov;14(3):285–91.

23. Pulst SM, Nechiporuk A, Nechiporuk T, Gispert S, Chen XN, Lopes-Cendes I, et al. Moderate expansion of a normally biallelic trinucleotide repeat in spinocerebellar ataxia type 2. Nat Genet. 1996 Nov;14(3):269–76.

24. Sanpei K, Takano H, Igarashi S, Sato T, Oyake M, Sasaki H, et al. Identification of the spinocerebellar ataxia type 2 gene using a direct identification of repeat expansion and cloning technique, DIRECT. Nat Genet. 1996 Nov;14(3):277–84.

25. Meierhofer D, Halbach M, Şen NE, Gispert S, Auburger G. Ataxin-2 (Atxn2)-Knock-Out Mice Show Branched Chain Amino Acids and Fatty Acids Pathway Alterations. Mol Cell Proteomics. 2016 May;15(5):1728–39.

26. Sen NE, Drost J, Gispert S, Torres-Odio S, Damrath E, Klinkenberg M, et al. Search for SCA2 blood RNA biomarkers highlights Ataxin-2 as strong modifier of the mitochondrial factor PINK1 levels. Neurobiol Dis. 2016 Dec;96:115–26.

27. Bakthavachalu B, Huelsmeier J, Sudhakaran IP, Hillebrand J, Singh A, Petrauskas A, et al. RNP-Granule Assembly via Ataxin-2 Disordered Domains Is Required for Long-Term Memory and Neurodegeneration. Neuron. 2018 May 16;98(4):754-766.e4.

28. Misgeld T, Schwarz TL. Mitostasis in Neurons: Maintaining Mitochondria in an Extended Cellular Architecture. Neuron. 2017 Nov 1;96(3):651–66.

29. Nissanka N, Moraes CT. Mitochondrial DNA damage and reactive oxygen species in neurodegenerative disease. FEBS Lett. 2018 Mar;592(5):728–42.

30. van Dijken JP, Bauer J, Brambilla L, Duboc P, Francois JM, Gancedo C, et al. An interlaboratory comparison of physiological and genetic properties of four Saccharomyces cerevisiae strains. Enzyme and Microbial Technology. 2000 Jun;26(9–10):706–14.

31. Longtine MS, McKenzie A, Demarini DJ, Shah NG, Wach A, Brachat A, et al. Additional modules for versatile and economical PCR-based gene deletion and modification in Saccharomyces cerevisiae. Yeast. 1998 Jul;14(10):953–61.

32. Maccecchini ML, Rudin Y, Blobel G, Schatz G. Import of proteins into mitochondria: precursor forms of the extramitochondrially made F1-ATPase subunits in yeast. Proc Natl Acad Sci U S A. 1979 Jan;76(1):343–7.

